# FFA and OFA encode distinct types of face identity information

**DOI:** 10.1101/2020.05.12.090878

**Authors:** Maria Tsantani, Nikolaus Kriegeskorte, Katherine Storrs, Adrian Lloyd Williams, Carolyn McGettigan, Lúcia Garrido

**Affiliations:** Division of Psychology, Department of Life Sciences, Brunel University London, Uxbridge, UK; Department of Psychological Sciences, Birkbeck, University of London, UK; Zuckerman Mind Brain Behavior Institute, Columbia University, New York, USA; Department of Experimental Psychology, Justus Liebig University, Giessen, Germany; Speech Hearing and Phonetic Sciences, University College London, London, UK; Department of Psychology, City, University of London, London, UK

**Keywords:** representational similarity analysis, face identity, FFA, OFA

## Abstract

Faces of different people elicit distinct functional MRI (fMRI) patterns in several face-selective brain regions. Here we used representational similarity analysis to investigate what type of identity-distinguishing information is encoded in three face-selective regions: fusiform face area (FFA), occipital face area (OFA), and posterior superior temporal sulcus (pSTS). We used fMRI to measure brain activity patterns elicited by naturalistic videos of famous face identities, and compared their representational distances in each region with models of the differences between identities. Models included low-level to high-level image-computable properties and complex human-rated properties. We found that the FFA representation reflected perceived face similarity, social traits, and gender, and was well accounted for by the OpenFace model (deep neural network, trained to cluster faces by identity). The OFA encoded low-level image-based properties (pixel-wise and Gabor-jet dissimilarities). Our results suggest that, although FFA and OFA can both discriminate between identities, the FFA representation is further removed from the image, encoding higher-level perceptual and social face information.

## Introduction

The human brain contains several face-selective regions that consistently respond more to faces than other visual stimuli (Kanwisher et al., 1997; Pitcher et al., 2011; Rossion et al. 2012). Functional magnetic resonance imaging (fMRI) has revealed that some of these regions represent different face identities with distinct brain patterns. Specifically, studies using fMRI multivariate pattern analysis have shown that face identities can be distinguished based on their elicited response patterns in the fusiform face area (FFA), occipital face area (OFA), posterior superior temporal sulcus (pSTS), and anterior inferior temporal lobe (Nestor et al. 2011; Verosky et al., 2013; Goesaert & Op de Beeck, 2013; Anzellotti et al., 2014; Axelrod & Yovel, 2015; Zhang et al., 2016; Anzellotti & Caramazza, 2017; Guntupalli et al., 2017; Visconti di Oleggio Castello et al., 2017; Tsantani et al., 2019). But do these regions represent the same information and, if not, what information is explicitly encoded in each of these face-selective regions?

Behaviourally, we distinguish between different faces using the surface appearance of the face, the shape of face features, and their spacing or configuration (e.g. Rhodes, 1988; Calder et al., 2001; Yovel & Duchaine, 2006; Russell & Sinha, 2007; Russell et al., 2007; Tardif et al., 2019). In particular, Abudarham and Yovel (2016) recently showed that features such as lip thickness, hair colour, eye colour, eye shape, and eyebrow thickness were crucial in distinguishing between individuals (see also Abudarham et al., 2019). Additionally, we perceive a vast amount of socially-relevant information from faces that can be used to distinguish between different individuals, such as gender, age, ethnicity, social traits (Oosterhof & Todorov, 2008; Sutherland et al. 2013), and even relationships and social network position (Parkinson et al., 2014; 2017). Therefore, if the response patterns in a certain brain region distinguish between two individuals, that region could be representing any one—or a combination of—these dimensions.

Like several other studies (see above), Goesaert and Op de Beeck (2013) demonstrated that the FFA, OFA, and a face-selective region in the anterior inferior temporal lobe could all decode between different face identities based on fMRI response patterns. Importantly, the authors further tested what type of face information was *encoded* in these different regions. The authors found that all three regions could distinguish between faces using both configural and featural face information, and therefore all regions seemed to represent similar information. Goesaert and Op de Beeck (2013) also showed that representational distances between different faces in face-selective regions did not correlate with low-level pixel-based information. This study however, used one single image for each person’s face, making it difficult to disentangle whether representations in a certain brain region are related to identity *per se* or related to the specific images used.

To determine whether brain response patterns represent face identity *per se*, others investigated whether those patterns could not only decode between faces of different people, but also show generalisation across different images of the same person’s face. Anzellotti et al (2014) showed that classifiers trained to decode face identities in the FFA, OFA, anterior temporal lobe, and pSTS (later analysed in Anzellotti and Caramazza, 2017) could also decode the same faces from novel viewpoints. Guntupalli et al (2017) additionally showed a hierarchical organisation of the functions of face-selective regions, with the OFA decoding viewpoint of face independently of the face identity, the anterior inferior temporal lobe (and a region in the inferior frontal cortex) decoding face identity independently of the viewpoint, and the FFA decoding both viewpoint and identity information. In contrast, Grossman et al (2019) have recently shown that representational distances between different face identities (computed from brain response patterns recorded from implanted electrodes) were very similar across the OFA and the FFA (in the left hemisphere). Crucially, the representational geometries in both regions were associated with differences in image-level descriptions computed from a deep neural network (VGG-Face), which were not generalisable across different viewpoints of the same person’s face. These results thus suggest that the OFA and FFA both represent complex configurations of image-based information and not face identity *per se*. Some studies have also shown that even lower-level stimulus-based properties of face images, such as those computed by Gabor filters, explain significant variance in the representational geometries in the FFA (Carlin & Kriegeskorte, 2017) as well as OFA and pSTS (Weibert et al., 2018). On the other hand, other studies have shown that more high-level information, such as biographical information and social context, affects the similarity of response patterns to different faces in the FFA (Verosky et al., 2013; Collins et al., 2016).

There is thus mixed evidence regarding whether different face-selective regions rely on similar or distinct information to distinguish between face identities, and what type of information may be encoded in different regions. In the present study, we used representational similarity analysis (RSA) (Kriegeskorte et al., 2008a; 2008b) to investigate what type of identity-distinguishing information is encoded in different face-selective regions. In our previous work (Tsantani et al., 2019), we showed that famous face-identities could be distinguished in the right FFA, OFA, and pSTS based on their elicited fMRI response patterns. Here, for the same set of famous identities and using the same data as in Tsantani et al (2019), we compared the representational distances between identity-elicited fMRI patterns in these regions with diverse candidate models of face properties that could potentially be used to distinguish between identities.

Importantly, we used multiple naturalistically varying videos for each identity that varied freely in terms of viewpoint, lighting, head motion, and general appearance. In addition, our representational distances were cross-validated across different videos, in order to deconfound identity from incidental image properties. By using a large, diverse set of candidate models, based on image properties of the stimuli (*image-computable models*) and on human-rated properties (*perceived-property models*), we were able to determine what types of identity-distinguishing information are encoded in different face-selective regions.

## Results

We tested 30 participants in an fMRI experiment, in which they were presented with faces of 12 famous people (same fMRI data as in Tsantani et al., 2019), and in a separate behavioural experiment, in which participants rated the faces of the same people on perceived similarity and social traits (Figure 1). We then computed representational dissimilarity matrices (RDMs) showing the representational distances between the brain response patterns elicited by the face identities in the face-selective right FFA, OFA, and pSTS (see Methods). The distance measure that we used to compute the RDMs was the linear discriminant contrast (LDC), which is a crossvalidated estimate of the Mahalanobis distance (Walther et al., 2016). The mean LDC across each RDM showed that response patterns to different face identities were discriminable in all three regions (Tsantani et al., 2019). To investigate the informational content of brain representations of the face identities in each face-selective region, we used RSA (Kriegeskorte et al., 2008a; 2008b) to compare the brain RDMs with a diverse set of candidate model RDMs (Figure 2). We used candidate models based on the physical properties of the stimuli (*image-computable models*), including low-level stimulus properties (based on Pixel-wise, GIST (Oliva & Torralba, 2001) and Gabor-jet (Biederman & Kalocsai, 1997) dissimilarities) and higher-level image-computable descriptions obtained from a deep neural network trained to cluster faces according to identity (OpenFace; Amos et al., 2016) (see Methods and Supplementary Information 1). Additionally, we used candidate models based on perceived higher-level properties (*perceived-property models*), including Gender and participants’ ratings of the face identities on Perceived Similarity and Social traits (Trustworthiness, Dominance, Attractiveness, Valence, and Social Traits (All) — which corresponds to all traits combined) in a behavioural experiment.

**Figure 1.**
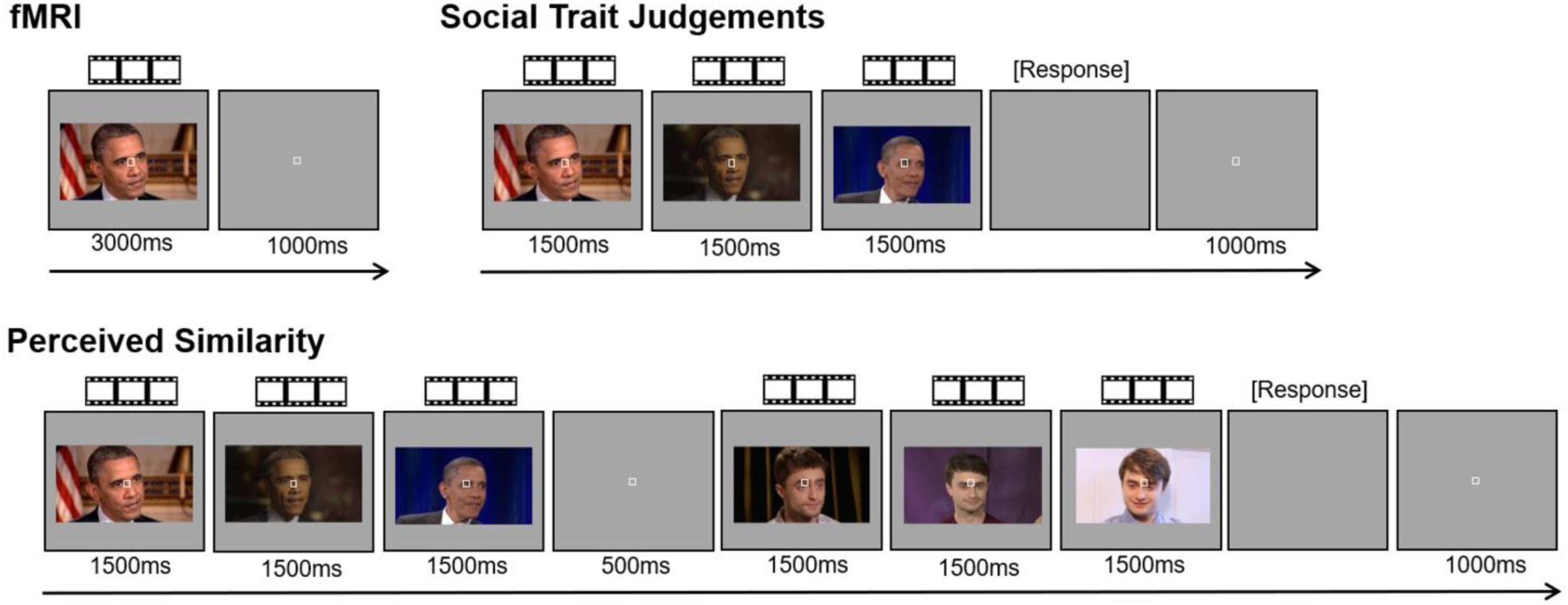
Examples of face trials in the fMRI and behavioural experiments. All experiments presented the same videos of moving, non-speaking, faces of 12 famous people. For each famous person, we presented six naturalistically varying videos of their face. In each fMRI trial, participants viewed a single face video. In each trial of the Social Trait Judgements Tasks (separate tasks for Trustworthiness, Dominance, Attractiveness, and Valence), participants viewed three videos of the face of the same identity and judged the intensity of the target trait (on a scale from 1 to 7). In each trial of the Perceived Similarity Task, participants viewed three videos of one identity followed by three videos of a different identity and rated their visual similarity (from 1 to 7). Face videos were presented for their full duration of 3000ms in the fMRI experiment, whereas only the first 1500ms were presented in the behavioural experiments.

**Figure 2.**
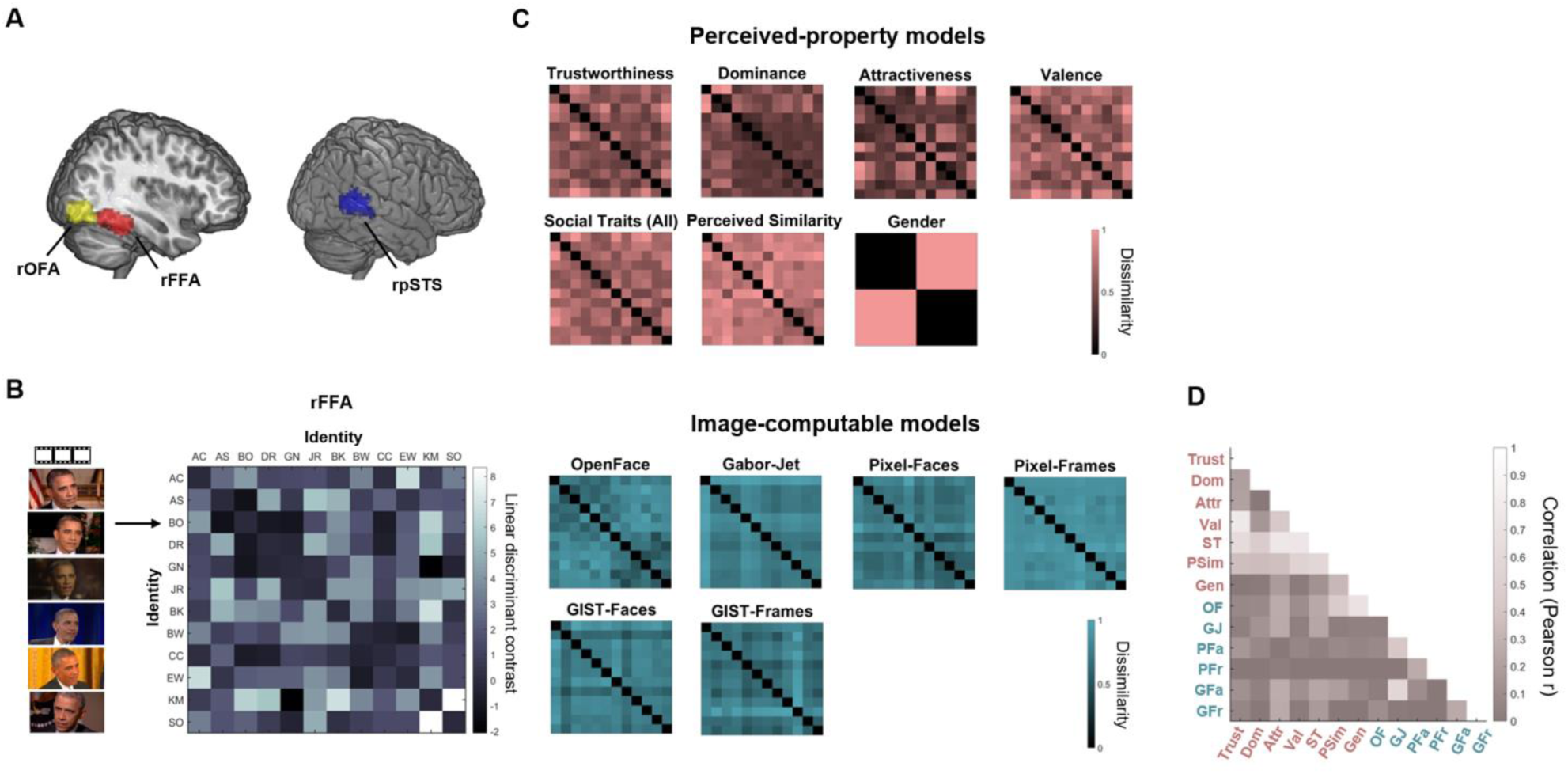
Brain and model representational dissimilarity matrices (RDMs). **A: Location in MNI space of the three face-selective regions localised in our participants**: OFA (occipital face area), FFA (fusiform face area), and pSTS (posterior superior temporal sulcus; all regions in the right hemisphere). These probabilistic maps were created for illustration purposes (in our analyses, we only used subject-specific regions of interest (ROIs)) and show all voxels that were present in at least 20% of participants. **B: Example brain representational dissimilarity matrix (RDM) for the right FFA**. For each ROI and each participant, we computed RDMs showing the dissimilarity of the brain response patterns between all pairs of identities. Each row and column represent one identity, and response patterns are based on all six presented videos of that identity. Each cell shows the linear discriminant contrast distance between the response patterns of two identities (higher values indicate higher dissimilarity), crossvalidated across runs presenting different videos of the face of each identity. The matrix is symmetric around a diagonal of zeros. **C: Model RDMs for *image-computable properties* (blue) and *perceived properties* (pink)**. These models are in the same format as the brain RDMs and show the dissimilarity between two identities on each property (see Methods). *Image-computable models* include a neural network trained to distinguish between face identities (OpenFace), a Gabor-Jet model, Pixel Dissimilarity (both for faces only — Pixel-Faces, and the whole frames — Pixel-Frames), and a GIST Descriptor model (both for faces — GIST-Faces, and the whole frames — GIST-Frames). *Perceived-property models* include perceived social traits (Trustworthiness, Dominance, Attractiveness, Valence, Social Traits (All)), Perceived Similarity, and Gender. Models based on participant ratings were averaged across participants. All models were built based on multiple images (image-computable models) or videos (perceived-property models) of the face of each identity. For visualisation purposes, all model RDMs were scaled to a range between zero (no dissimilarity) and one (maximum dissimilarity). **D: Correlations (Pearson) between the different model RDMs.** The different candidate models were compared with each other using Pearson correlation.

### Individual model analysis

In our main analysis, we computed Pearson’s correlations between RDMs in the right FFA, OFA, and pSTS, and each candidate model RDM. Correlations were computed for each individual participant, and then correlations across participants for each model were compared against zero using two-sided one-sample Wilcoxon signed-rank tests. For each ROI and each model that showed significant correlations with participants’ brain RDMs, we report below the mean correlation across participants, and the Z statistic and p-value obtained from the signed-rank test, corrected for multiple comparisons using FDR correction. Full results are presented in Table S2-1, and individual-subject correlations are presented in Figure S2-1 (Supplementary Information 2). We also compared the correlations across all pairs of models using two-sided Wilcoxon signed-rank tests.

Brain RDMs in the right FFA had the highest mean correlation with the Perceived Similarity model (mean *r* = .11, *Z* = 3.69, *p* = .0002), followed by perceived Social Traits (All) (mean *r* = .10, *Z* = 2.71, *p* = .0067), the image-computable neural network OpenFace (mean *r* = .10, *Z* = 3.46, *p* = .0005), perceived Attractiveness (*mean r* = .09, *Z* = 2.69, *p* = .0072), Gender (mean *r* = .09, *Z* = 3.30, *p* = .0010), and Valence (mean *r* = .06, *Z* = 2.39, *p* = .0168) (Figure 3A). We estimated the lower bound of the noise ceiling as the mean correlation between each participant’s FFA RDM and the average of all other participants’ FFA RDMs (Nili et al., 2014). This estimates the non-noise variance in the data, and is not overfit to the present data. None of the mean correlations reached the lower bound of the noise ceiling for the FFA (*r* = .14) — this suggests that there could be models outside those tested here that would better explain the representational distances in FFA. Pairwise comparisons showed no significant differences between the correlations of any pairs of models (all p>.0041; no significant results after FDR correction).

**Figure 3:**
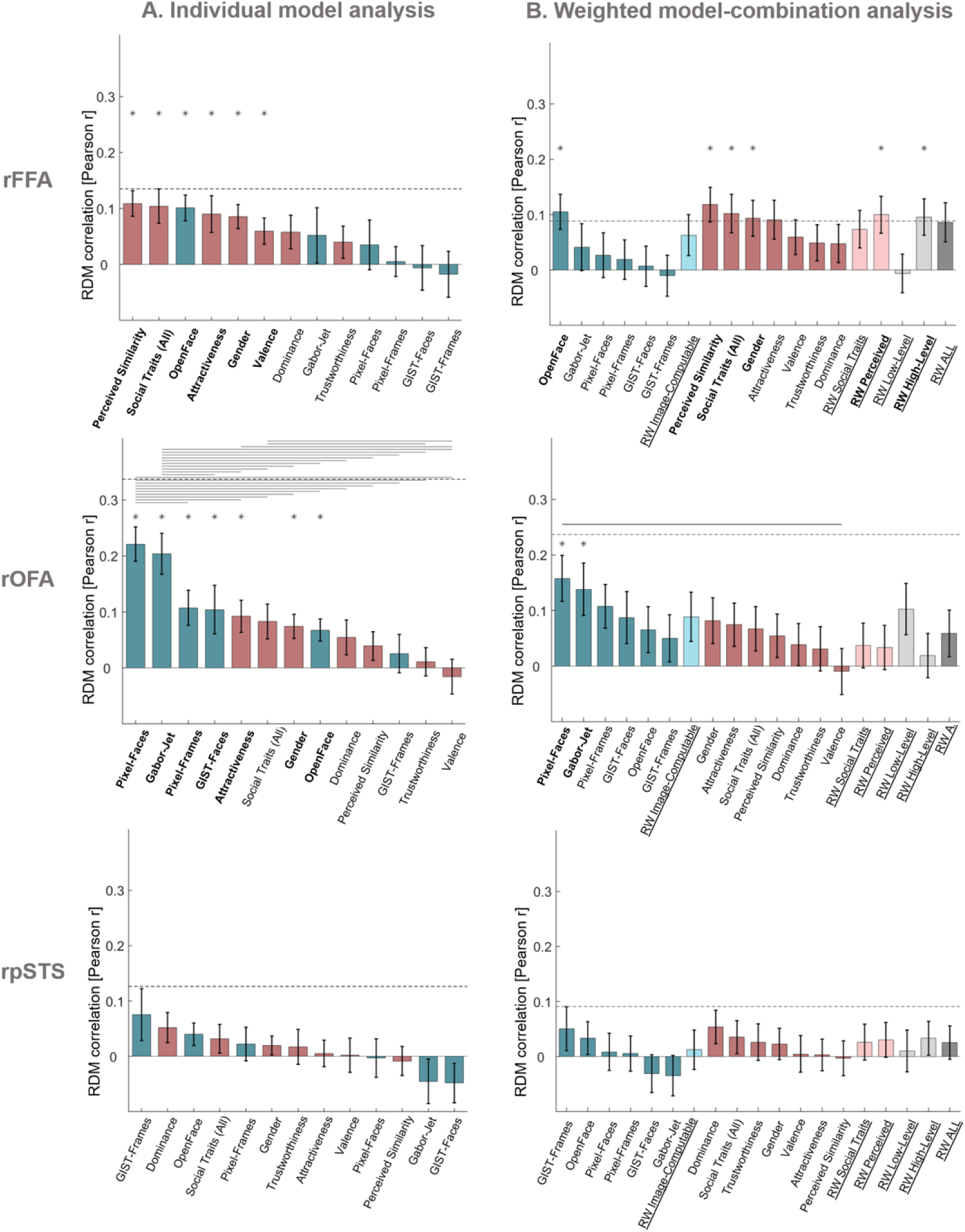
FFA and OFA show distinct representational profiles of face identity information. **A: Similarity (Pearson correlations) between brain RDMs (in FFA, OFA, and pSTS) and each of the individual candidate models**. Bars show mean correlations across participants and error bars show standard error. Correlations with image-computable models are in blue and with perceived-property models are in pink. Horizontal dashed lines show the lower bound of the noise ceiling. An asterisk above a bar and the name of the model in bold indicate that correlations with that model were significantly higher than zero. Correlations with individual models are sorted from highest to lowest. Horizontal lines above bars show significant differences between the correlations of the two end points (FDR corrected for multiple comparisons). **B: Similarity (Pearson correlations) between brain RDMs (in FFA, OFA, and pSTS) and each of the candidate models in the weighted representational modelling analysis**. Bars show mean correlations and error bars show standard error across 1,000 bootstrap samples. Horizontal dashed lines show the lower bound of the noise ceiling, averaged across bootstrap samples. An asterisk above a bar and the name of the model in bold indicate that correlations with that model were significantly higher than zero. Correlations with individual models are blocked by type of model (image-computable models followed by perceived-property models) and sorted from highest to lowest. RW shows the combined and reweighted models and appears in light blue for models that combine image-computable properties, in light pink for models that combine Perceived properties, and in grey for models that combine both types of properties. None of the combined models outperformed individual models. The results of both analyses show that in the FFA, the models that explained most of the variance are related to high-level properties, such as perceived properties of the stimuli and the image-computable OpenFace model of face recognition. In contrast, brain RDMs in OFA correlated mainly with low-level image-computable properties such as pixel dissimilarity and the Gabor-Jet model. No significant correlations were found in pSTS.

In contrast with the FFA, the brain RDMs in the right OFA had the highest mean correlations with low-level image-computable models. The highest mean correlation was observed with the Pixel-Faces model (mean *r* = .22, *Z* = 4.36, *p* < .0001) (Figure 3A), followed by the Gabor-Jet (mean *r* = .20, *Z* = 3.97, *p* < .0001), Pixel-Frames (mean *r* = .11, *Z* = 3.02, *p* = .0026), GIST-Faces (mean *r* = .10, *Z* = 2.22, *p* = .0267), perceived Attractiveness (mean *r* = .09, *Z* = 2.84, *p* = .0045), Gender (mean *r* = .07, *Z* = 2.76, *p* = .0058), and the OpenFace model (mean *r* = .07, *Z* = 2.95, *p* = .0032). None of the mean correlations reached the lower bound of the noise ceiling (*r* = .34). Pairwise comparisons between model correlations revealed that the Pixel-Faces model had significantly higher correlations with the OFA RDMs than all other models (all *p* < .0058, FDR corrected), except for the Gabor-Jet model and the GIST-Faces model. The Gabor-Jet model also had significantly higher correlations with the brain RDMs in OFA than all other models (all *p* < .0058, FDR corrected), except the Pixel-Faces and Pixel-Frames models. Perceived Attractiveness had significantly higher correlations with the OFA RDMs than perceived Valence (*p* = .0051), and Social traits (All) was significantly higher than Trustworthiness and Valence (both *p* < .0018).

Finally, we investigated which model best explained the variance in representational distances in the right pSTS. We found no significant correlations between any of the candidate models and the brain RDMs in this region (all p > .0333; no significant results after FDR correction) (Figure 3A). None of the models reached the lower bound of the noise ceiling (*r* = .13), and there were no significant differences between models (all *p* > .0140; no significant results after FDR correction).

These results show a clear distinction between the types of models that were associated with the representational geometries of face-identities in the FFA and OFA. Representational distances of face identities in the FFA were most associated with high-level perceived similarity, gender, and social traits, as well as a high-level model of image-computable properties (OpenFace), whereas representations in OFA were most associated with low-level image-computable properties. To further test the hypothesis that the FFA and OFA encode different information about face identities, we computed correlations between each participant’s brain RDM for each ROI, and the mean brain RDM for all other participants in the same ROI, and in each of the other ROIs. Our results showed that for all three ROIs, participants’ brain RDMs were on average more similar to other participants’ RDMs for the same ROI than to other participants’ RDMs of different ROIs, suggesting that there are robust region-specific computations in each ROI that generalise across individuals (Supplementary Information 3). We additionally showed that we obtained similar results to those in Figure 3A when using other similarity measures between RDMs (Spearman correlation, Kendall tau-a), demonstrating that these results are not dependent on using Pearson correlation (Supplementary Information 4). Finally, we conducted an additional control analysis using brain RDMs in the same ROIs but built from response patterns to voices of the same individuals, instead of brain responses to faces. There were no significant correlations between any of the model RDMs for faces and brain RDMs for voices (Supplementary Information 5), demonstrating that the above results for FFA and OFA are specific to visual stimuli (faces). To conclude, we find that the structure of the model correlations is reliable and is systematically different between the FFA and OFA.

### Weighted model-combination analysis

Although our models accounted for a large portion of the explainable variance (based on the noise ceiling) in brain representations in the right FFA and OFA, none of the mean correlations reached the lower bound of the noise ceiling. It could be that each individual model captured only a portion of the information represented in each brain region, in which case we may be able to fully explain the brain representations by combining multiple models. We thus used weighted representational modelling (Khaligh-Razavi & Kriegeskorte, 2014; Jozwik et et al., 2016; Jozwik et al., 2017) to combine sets of models into weighted combinations via crossvalidated fitting on the human data, and to investigate if these combined models resulted in better predictions of the brain dissimilarities in each brain region (see Methods). We considered six different combined models: *Image-computable* properties (OpenFace, GIST, GaborJet, and Pixel), *Social Traits* (comprising a weighted combination of the Trustworthiness, Dominance, Attractiveness, and Valence properties), *Perceived* properties (Trustworthiness, Dominance, Attractiveness, Valence, Perceived Similarity, and Gender), *Low-Level* properties (GIST, GaborJet, and Pixel), *High-Level* properties (Trustworthiness, Dominance, Attractiveness, Valence, Perceived Similarity, Gender, and OpenFace), and *All properties*.

We used linear non-negative least squares regression to estimate a weight for each component of each combined model. We fitted the weights and tested the performance of the reweighted (combined) model on non-overlapping groups of both participants and stimulus conditions within a cross-validation procedure, and used bootstrapping to estimate the distribution of the combined model’s performance (Storrs et al., 2020). Figure 3B shows the results of this analysis. P-values were corrected for multiple comparisons using Bonferroni correction. For the FFA, the combined models for Perceived properties and High-Level properties had the highest mean correlations with the brain RDMs, and the individual-subject correlations were significantly above zero. For the OFA, the combined model of all Low-Level properties and that of all image-computable properties had the highest mean correlations with the brain RDMs, although the individual-subject correlations were not significantly above zero after correcting for multiple comparisons. Importantly, however, none of the combined models performed better than the best of the individual models (see full results in Supplementary Information 6). Instead, the models with best performance in the previous (main) analysis also showed the highest correlations in this analysis. These results suggest that the models that best explained representational distances in each face-selective region share overlapping variance, given that combining them did not improve model performance. Lastly, replicating the findings of the previous analysis using more stringent statistical methods (crossvalidation across stimuli and participants) provides further evidence of a reliable pattern of model correlations in FFA and OFA that reveals a distinction between the type of information encoded in these two regions.

## Discussion

We aimed to investigate what information is explicitly encoded in the face-selective right FFA, OFA, and pSTS. We extracted fMRI patterns elicited by famous face identities in these regions, and computed face identity RDMs which showed that face identities could be distinguished based on their elicited response patterns in all three regions. Using RSA, we compared the brain RDMs for the FFA, OFA, and pSTS with multiple model RDMs ranging from low-level image-computable properties (pixel-wise, GIST, and Gabor-jet dissimilarities), through higher-level image-computable descriptions (OpenFace deep neural network, trained to cluster faces by identity), to complex human-rated face properties (perceived visual similarity, social traits, and gender). We found that the FFA and rOFA encode face identities in a different manner, suggesting distinct representations in these two regions. The representational geometries of face identities in the FFA were most associated with high-level properties, such as perceived visual similarity, social traits, gender, and high-level image features extracted with a deep neural network (OpenFace; Amos et al., 2016). In contrast, the representational geometries of faces in the right OFA were most associated with low-level image-based properties, such as pixel similarity and features extracted with Gabor filters that simulate functioning of early visual cortex. While previous studies had shown that low-level properties of images extracted with Gabor filters were associated with representational distances of faces in right FFA (Carlin & Kriegeskorte, 2017; Weibert et al, 2018), our results suggest that representations in right FFA use more complex combinations of stimulus-based features and relate to higher-level perceived and social properties. These results inform existing neurocognitive models of face processing (Haxby et al., 2000; Duchaine & Yovel, 2015) by shedding light on the much-debated computations of face-responsive regions, and providing new evidence to support a hierarchical organisation of these regions from the processing of low-level image-computable properties in the OFA to higher-level visual features and social information in the FFA.

Our initial prediction was that by combining and reweighting different candidate models, we would be better able to explain the brain RDMs. However, we did not find evidence for this in any of our face-selective ROIs. These results suggest that, when more than one model was significantly correlated with the brain RDMs for a certain brain region, they tended to explain overlapping variance in the brain RDMs. For example, while Perceived Similarity and OpenFace both explained the representational geometries in right FFA, their combination did not explain more variance than each model individually. However, our pattern of results suggests a clear distinction between the *types* of models that are associated with representations in the FFA and OFA, with higher-level properties explaining more variance in the FFA, and lower-level image-based properties explaining more variance in the OFA.

One crucial aspect of our study is that we used naturalistically varying video stimuli and multiple tokens (depictions) for each identity. Brain RDMs were built by cross-validating the response patterns across runs featuring different videos of the face of each identity, and behavioural models were based on averages of ratings of multiple videos for each identity. Image-based models were built by calculating dissimilarities between image frames taken from multiple videos of the face of each identity, and then computing the mean dissimilarity across different image pairs featuring the same identity pair. Behavioral studies have demonstrated that participants make more mistakes in “telling together” (i.e. grouping multiple images of the same identity, which is different process from “telling apart”, or distinguishing, between different identities) different photos of the same person when those photos were taken with different cameras, on different days, or with different lighting conditions, compared to when photos were taken on the same day and with the same camera (Bruce et al, 1999, Jenkins et al, 2011). Most previous fMRI studies, however, used very visually similar images, or even just a single image, for each identity, making it difficult to determine whether a brain region represents different *face images* or different *face identities*. Here, by having multiple videos/tokens for each person we can be more confident that we are capturing representations of specific identities rather than specific stimuli.

Related to the previous point, Abudarham and Yovel (2016) have recently shown that humans are more sensitive in perceiving changes in some face features (such as lip-thickness, hair, eye colour, eye shape, and eyebrow thickness) compared to others (such as mouth size, eye distance, face proportion, skin color). Changes in the former type of features (a.k.a. critical features) are perceived as changes in identity and those features tend to be invariant for different tokens of the same identity. Interestingly, Abudarham et al (2019) showed that the OpenFace algorithm that we used in the present study also seemed to be capturing those same critical features. Given our results in right FFA, it would be interesting to see whether representations in this region can also distinguish between the processing of the critical and non-critical face features as described by Abudarham and colleagues (2016; 2019).

Grossman and colleagues (2019) have also recently shown that representations in the FFA relate to image-computable descriptors from a deep neural network. There are two main differences, however, between our results and those of Grossman et al (2019). First, Grossman et al (2019) found similar representational geometries across all face-selective ventral temporal cortex, and no differentiation between OFA and FFA. One possible reason for this difference is that the authors were only able to define OFA and FFA in the left hemisphere, whereas our face-selective regions were defined in the right hemisphere. Face-selective regions are more consistent and larger in the right hemisphere (e.g. Rossion et al, 2012). A second main difference between our results and those of Grossman et al (2019) is that the deep neural network that we used here showed high generalisation across different images of the same person. OpenFace (Amos et al., 2016) was trained specifically to group together images of the same person and distinguish images of different people, and it performed very well in doing this in our set of stimuli (see Supplementary Information 1), where it showed high generalisation across very variable pictures of the same person. This was not the case with the VGG-Face network used by Grossman et al (2019). Future studies should focus on describing and comparing the image-level descriptions of different types of neural networks.

We note that the lower bounds on the noise ceiling in our analyses were consistently quite low, especially for FFA and pSTS. However, these values are similar to the lower bounds of the noise ceiling in other studies using RSA (e.g. Carlin & Kriegeskorte, 2017; Jozwik et al., 2016; Thornton & Mitchell, 2017; 2018). The low noise ceilings in our study likely reflect the fact that the differences between brain-activity patterns associated with faces of different people are small compared to the differences between patterns associated with different visual categories (e.g. faces and places). Moreover, we used identity-based rather than image-based patterns (by crossvalidating across runs presenting different tokens for each identity), and this is likely to have introduced additional variability to the pattern estimates. It is also possible that we needed more data per participant, and future studies should consider ways to increase the amount of explainable variance.

Considering that we defined the noise ceiling as inter-subject reliability (correlations between RDMs of different participants), an alternative interpretation of the low noise ceilings could be that there were substantial individual differences in face identity brain representations. It is possible that brain RDMs are reliable within the same individual but less so across individuals. If this were the case, each person’s brain RDMs may be well explained by a uniquely weighted combination of candidate models, but no set of weightings would perform well for all participants. In supplementary analyses we tested whether it is possible that brain RDMs are more reliable within the same individual (across different scanning sessions) than across individuals, and whether behavioural models based on participants’ own ratings (for perceived similarity and perceived social traits) may be better predictors of their own brain representational distances compared to models based on ratings averaged across participants (Supplementary Information 7). Our results, however, showed that intra-subject reliability of brain RDMs was in fact lower than the inter-subject reliability, and that using the participant’s own individual behavioural models did not improve correlations with their brain RDMs.

None of the models that we considered here explained the representational geometry of responses in the face-selective right pSTS. It is likely that the pSTS as defined in the present study contains overlapping and interspersed groups of voxels that respond to faces only, voices only, or both faces and voices (Beauchamp et al., 2004) that make the overlapping representational geometry difficult to explain. On the other hand, it is possible that the pSTS represents information about people that we did not consider here, such as idiosyncratic facial movements (Yovel & O’Toole, 2016), emotional and mental states (Thornton et al., 2019), social distance or network position (Parkinson et al., 2014; 2017), or type of social interactions (Walbrin & Koldewyn, 2019). Future studies may need to explore an even richer set of social, perceptual, and stimulus-based models to better characterise responses in the pSTS.

To conclude, our study highlights the importance of using multiple and diverse representational models to characterise how face identities are represented in different face-selective regions. Although similar levels of identity decodability were observed in both OFA and FFA (Tsantani et al., 2019), the information explicitly encoded in these two regions is in fact distinct, suggesting that the two regions serve quite different computational roles. Future work attempting to define the computations of cortical regions that appear to serve the same function (e.g. discriminating between identities) would benefit from comparing representations in those regions with multiple and diverse candidate models to reveal the type of information that is encoded.

## Materials and Methods

This study involved an fMRI component, in which we measured brain representations of faces and voices, and a behavioural component, in which we collected ratings of the same faces and voices on social traits and perceived similarity. The fMRI part corresponds to the same experiment and data described in Tsantani et al (2019) and the behavioural part is reported here for the first time. In the present study, we analysed the data related to faces only.

### Participants

We recruited thirty-one healthy right-handed adult participants to take part in two fMRI sessions and a behavioural session (all on separate days, resulting in at least six hours of testing per participant). To ensure adequate exposure to our stimulus set of famous people, participants were required to be native English speakers between 18 and 30 years of age, and to have been resident in the UK for at least 10 years. We also independently verified that all participants knew the famous people used in the experiment (please see Tsantani et al., 2019). Participants were recruited at Royal Holloway, University of London, and Brunel University London. One participant was excluded due to excessive head movement in the scanner. The final sample consisted of 30 participants (eight men) with a mean age of 21.2 years (SD=2.37, range=19-27). Participants reported normal or corrected-to-normal vision and normal hearing, provided written informed consent, and were reimbursed for their participation. The study was approved by the Ethics Committee of Brunel University London.

### Stimuli

The same stimuli were used in the fMRI and behavioural testing, and consisted of videos of the faces and sound recordings of 12 famous individuals, including actors, comedians, TV personalities, pop stars and politicians: Alan Carr, Daniel Radcliffe, Emma Watson, Arnold Schwarzenegger, Sharon Osbourne, Graham Norton, Beyonce Knowles, Barbara Windsor, Kylie Minogue, Barack Obama, Jonathan Ross, and Cheryl Cole. These individuals were selected based on pilot studies that showed that participants (aged between 18 and 30 and living in the UK) could recognise them easily from their faces and voices.

For each identity, six silent, non-speaking video clips of their moving face were obtained from videos on YouTube. The six tokens were obtained from different original videos. In total, we obtained 72 face stimuli. Face videos were selected so that the background did not provide any cues to the identity of the person. The face videos were primarily front-facing and did not feature any speech but were otherwise unconstrained in terms of facial motion. Head movements included nodding, smiling, and rotating the head. Videos were edited so that they were three seconds long, 640 x 360 pixels, and centred on the bridge of the nose, using Final Cut Pro X (Apple, Inc.).

For purposes not related to this study, we also presented 72 voice stimuli, which consisted of recordings of the voices of the same 12 famous individuals (6 clips per identity) obtained from videos on YouTube. Speech clips were selected so that the speech content, which was different for every recording, did not reveal the identity of the speaker. Recordings were edited so that they contained three seconds of speech after removing long periods of silence using Audacity® 2.0.5 recording and editing software (RRID:SCR_007198). The recordings were converted to mono with a sampling rate of 44100, low-pass filtered at 10KHz, and root-mean-square (RMS) normalised using Praat (version 5.3.80; Boersma and Weenink 2014; www.praat.org).

Participants were familiarised with all stimuli via one exposure to each clip immediately before the first scanning session.

### MRI data acquisition and preprocessing

Participants completed two MRI sessions: in each session, participants completed a structural scan, three runs of the main experiment, and functional localiser scans (for face and voice areas, but below we only describe the localiser of face-selective regions). Participants were scanned using a 3.0 Tesla Tim Trio MRI scanner (Siemens, Erlangen) with a 32-channel head coil. Scanning took place at the Combined Universities Brain Imaging Centre (CUBIC) at Royal Holloway, University of London. We acquired whole-brain T1-weighted anatomical scans using magnetization-prepared rapid acquisition gradient echo (MPRAGE) [1.0 × 1.0 in-plane resolution; slice thickness, 1.0mm; 176 axial interleaved slices; PAT, Factor 2; PAT mode, GRAPPA (GeneRalized Autocalibrating Partially Parallel Acquisitions); repetition time (TR), 1900ms; echo time (TE), 3.03ms; flip angle, 11°; matrix, 256×256; field of view (FOV), 256mm].

For the functional runs, we acquired T2*-weighted functional scans using echo-planar imaging (EPI) [3.0 × 3.0 in-plane resolution; slice thickness, 3.0mm; PAT, Factor 2; PAT mode, GRAPPA; 34 sequential (descending) slices; repetition time (TR), 2000ms; echo time (TE), 30ms; flip angle, 78°; matrix, 64×64; field of view (FOV), 192mm]. Slices were positioned at an oblique angle to cover the entire brain except for the most dorsal part of the parietal cortex. Each run of the main experiment comprised 293 brain volumes, and each run of the face localizer had 227 brain volumes.

Functional images were pre-processed used Statistical Parametric Mapping (SPM12; Wellcome Department of Imaging Science, London, UK; RRID:SCR_007037; http://www.fil.ion.ucl.ac.uk/spm) operating in Matlab (version R2013b; MathWorks; RRID:SCR_001622). The first three EPI images in each run served as dummy scans to allow for T1-equilibration effects and were discarded prior to pre-processing. Data from each of the two scanning sessions, which took place on different days, were first pre-processed independently with the following steps for each session. Images within each brain volume were slice-time corrected using the middle slice as a reference, and were then realigned to correct for head movements using the first image as a reference. The participants’ structural image in native space was coregistered to the realigned mean functional image, and was segmented into grey matter, white matter, and cerebrospinal fluid. Functional images from the main experimental runs were not smoothed, whereas images from the localiser runs were smoothed with a 4-mm Gaussian kernel (full width at half maximum). To align the functional images from the two scanning sessions, the structural image from the first session was used as a template, and the structural image from the second session was coregistered to this template; we then applied the resulting transformation to all the functional images from the second session.

### Functional localisers and definition of regions of interest

Face-selective regions were defined using a dynamic face localiser that presented famous and non-famous faces, along with a control condition consisting of objects and scenes. The stimuli were silent, non-speaking videos of moving faces, and silent videos of objects and scenes, presented in an event-related design. Participants completed between one and two runs of the localiser across the two scanning sessions. The localiser presented different stimuli in each of two runs. For full details of the localiser please see Tsantani et al (2019).

Functional regions of interest (ROIs) were defined using the Group-Constrained Subject-Specific method (Fedorenko et al., 2010; Julian et al., 2012), which has the advantage of being reproducible and reducing experimenter bias by providing an objective means of defining ROI boundaries. Briefly, subject-specific ROIs were defined by intersecting subject-specific localiser contrast images with group-level masks for each ROI obtained from an independent dataset. In this study, we obtained group masks of face-selective regions (right fusiform face area (rFFA), the right occipital face area (rOFA), and the right posterior superior temporal sulcus (rpSTS)) from a separate group of participants who completed the same localiser (for details see Tsantani et al., 2019). We focused on face-selective regions from the right hemisphere because they have been shown to be more consistent and larger compared to the left hemisphere (e.g. Rossion et al, 2012). Our masks are publicly available at https://doi.org/10.17633/rd.brunel.6429200.v1.

Contrast images were defined for each individual participant. Face-selectivity was defined by contrasting activation to faces versus non-face stimuli using *t*-tests. We then intersected these subject-specific contrasts with the group masks, and extracted all significantly activated voxels at *p*<.001 (uncorrected) that fell within the boundaries of each mask. In cases where the resulting ROI included fewer than 30 voxels, the threshold was lowered to *p* <. 01 or *p* < .05. ROIs which included fewer than 30 voxels at the lowest threshold were not included, and this occurred for the rFFA in two participants and for the rOFA in one participant. For full details of size and location of all ROIs, please see Tsantani et al (2019).

## Experimental design

### Main experimental fMRI runs

In the main experimental runs, face stimuli were presented intermixed with voice stimuli within each run in an event-related design. The experiment was programmed using the Psychophysics Toolbox (version 3; RRID:SCR_002881; Brainard 1997; Pelli 1997) in Matlab and was displayed through a computer interface inside the scanner. Participants were instructed to fixate on a small square shape that was constantly present in the centre of the screen. From a distance of 85cm, visual stimuli subtended 20.83 × 12.27 degrees of visual angle on the 1024 × 768 pixel screen.

The six tokens of the face of each identity were evenly distributed across the three runs so that each run contained two unique videos of the face of each of the 12 identities (Figure 1). Each stimulus was presented twice, resulting in 96 experimental trials of faces and voices in each run. Stimuli were presented in a pseudorandom order that prohibited the succeeding repetition of the same stimulus and ensured that each identity could not be preceded or succeeded by another identity more than once within the same modality. Each trial presented a stimulus for 3000 ms and was followed by a 1000 ms ITI (Figure 1).

To maintain attention to stimulus identity in the scanner, participants performed an anomaly detection task in which they indicated via button press when they were presented with a famous face or voice that did not belong to one of the 12 famous individuals that they had been familiarised with prior to the experiment. Therefore, each run also included 12 randomly presented task trials (six faces & six voices). Finally, each run contained 36 randomly interspersed null fixation trials, resulting in a total of 144 trials in each run lasting around 10 minutes.

The three experimental runs that were completed in the first scanning session were repeated in the second session with the same stimuli, but in a new pseudorandom order. The task stimuli, however, were always novel for each run. The three runs, which had different tokens, were presented in counterbalanced order across participants in both sessions.

### Behavioural session

All participants completed a behavioural session in a laboratory, which took place on a separate day and always after the fMRI sessions had been completed. In this session, participants rated the same faces that they had been presented with in the scanner on perceived social traits and on perceived pairwise visual similarity. Participants also rated voices (the order of tasks was counterbalanced across modality), but these results are not presented here. All tasks and stimuli were presented using the Psychophysics Toolbox and Matlab.

#### Social Trait Judgement Tasks

In the social trait judgement tasks, participants were asked to make judgements about the perceived trustworthiness, dominance, attractiveness, and positive-negative valence of the face identities. There were four blocks, one for each judgement, and their order was counterbalanced across participants. Face stimuli were presented in the centre of the screen. In contrast to the fMRI runs, in which stimuli were presented for the full three seconds of their duration, here all stimuli were only presented for the first 1500 ms of their duration, to reduce testing time.

All blocks followed the same trial structure (Figure 1). In each trial, a face identity was presented with three videos — these were presented successively with no gap in between them (total of 4500 ms). Participants were then asked to rate how trustworthy/dominant/attractive/negative-positive the face was, and they were asked to base their judgement on all three videos of the face. The rating scale ranged from 1 (very untrustworthy/non-dominant/unattractive/negative) to 7 (very trustworthy/dominant/attractive/positive) and participants responded using the corresponding keys on the keyboard. There was a 1000ms ITI following the response.

Each identity was presented in two trials; one trial presented three face videos randomly selected from the six available, and the other trial presented the remaining three videos. This resulted in 24 trials in each block (12 identities × 2 presentations). Tokens within each trial were presented in a random order, and the trial order was also randomised. Trustworthiness was defined as ‘able to be relied on as honest and truthful’. Dominance was defined as ‘having power and influence over other people’. No definition was deemed necessary for valence or attractiveness. Participants were advised that there was no time limit to their responses and that they should follow their first judgment. The duration of each block was approximately 3 minutes.

#### Pairwise Visual Similarity Task

In the pairwise similarity task, participants rated the perceived visual similarity of pairs of face identities. Each of the 12 identities was paired with the other 11 identities creating 66 identity pairs. Each identity was presented by three videos, randomly selected from the six available videos. Each identity pair was presented in two trials, counterbalancing the presentation order of each identity in the pair. There were therefore 132 trials in each task (66 identity pairs × 2 presentations). The presentation order of the pairwise similarity tasks in relation to the social trait judgement tasks was also counterbalanced across participants.

Participants were instructed to rate the similarity between the visual appearance of the two face identities in each pair, focusing on the facial features. Participants were asked to rate how similar the two faces looked on a scale from 1 (very dissimilar) to 7 (very similar). Participants were advised that there was no time limit to their responses and that they should follow their first instinct. Participants were told to ignore similarities between people that were related to biographical or semantic information (e.g. if both identities were actors). Furthermore, to encourage participants to base their judgements on perceptual information, participants were advised to consider to what extent two identities could potentially be related to each other, i.e. be part of the same family, based on how they looked.

In each trial, participants were first presented with the three videos of the face of one identity (Figure 1). Following a 500ms fixation screen, they were presented with the three videos of the face of the second identity. Videos for each identity were presented successively with no gap in between. Each video was presented for 1500ms and there was a 1000ms ITI following the response. The presentation order of the trials was randomised. The duration of each task was approximately 30 minutes.

## Representational dissimilarity matrices (RDMs)

### Brain RDMs

Representational dissimilarity matrices (RDMs) showing the discriminability of the brain response patterns elicited by the 12 face identities (during the fMRI experimental runs) were created for each individual participant and for each ROI.

First, to obtain brain responses at each voxel for each of the 12 face identities, mass univariate time-series models were computed for each participant using a high-pass filter cutoff of 128 seconds and autoregressive AR(1) modelling to account for serial correlation. Regressors modelled the BOLD response at stimulus onset and were convolved with a canonical hemodynamic response function (HRF). We defined a model for each run separately, and for every possible pair of runs within a scanning session (by concatenating the two runs), to create data partitions for cross-validation (described below). Each model contained a regressor for the face of each of the 12 identities, which incorporated the different videos of their face (two per run) and the repetitions of those videos. The model also included regressors for each of the 12 voice identities, task trials, and the six motion parameters obtained during the image realignment preprocessing stage (included as regressors of no interest).

Second, within each ROI, we extracted the beta estimates at each voxel for each of the 12 face identities. This resulted in 12 vectors of beta values per ROI that described the response patterns (across voxels) elicited by the 12 face identities.

Third, these vectors of beta estimates were used to compute 12×12 Face RDMs in face-selective ROIs, in which each cell showed the distance between the response patterns of two identities (Figure 2B). RDMs were computed using the linear discriminant contrast (LDC), a cross-validated distance measure (Nili et al. 2014; Walther et al. 2016), which we implemented using in-house Matlab code and the RSA toolbox (Nili et al. 2014). Two RDMs were created for each ROI, one for each scanning session. Each RDM was computed using leave-one-run-out cross-validation across the three runs, which presented different stimuli for each identity. Therefore, RDMs showed the dissimilarities between face *identities*, rather than specific face videos. In each cross-validation fold, concatenated data from two runs formed partition A, and data from the left-out run formed partition B. For each pair or identities (e.g. ID1 and ID2), partition A was used to obtain a linear discriminant, which was then applied to partition B to test the degree to which ID1 and ID2 could be discriminated. Under the null hypothesis, LDC values are distributed around zero when two patterns cannot be discriminated. Values higher than zero indicate higher discriminability of the two response patterns (Walther et al. 2016).

The discriminability of face identities in each ROI was computed by calculating the mean LDC across all cells of each participant’s RDM, and comparing the mean LDC distances against zero (Tsantani et al., 2019).

Full details of this analysis are presented in Tsantani et al (2019) and the data to compute brain RDMs are available at https://doi.org/10.17633/rd.brunel.6429200.v1. Here, we used the RDMs for three face-selective regions (rFFA, rOFA, and rpSTS). All three of these regions showed significant discriminability of face identities.

### RDMs based on image-computable properties

We computed dissimilarities between the 12 face identities based on visual descriptions of their faces obtained using the models described below. We did not use the full videos as input to these models, but instead extracted one still frame from each face video used in the experiment (typically the first frame in which the full face was visible and the image was not blurred). Thus, we obtained six different images of the face of each identity, taken from the six different videos in which the identity was presented, resulting in 72 images in total.

#### OpenFace Model

The ‘OpenFace’ model RDM was computed from low-dimensional face representations obtained from OpenFace (Amos et al., 2016; http://cmusatyalab.github.io/openface/). Briefly, OpenFace uses a deep neural network that has been pre-trained (using 500,000 faces) to learn the best features or measurements that can group two pictures of the same identity together and distinguish them from a picture of a different identity. We used this pre-trained neural network to generate measurements for each of our face pictures and to compare these measurements between each pair of pictures. OpenFace first performs face-detection, identifies pre-specified landmarks, and does an affine transformation so that the eyes, nose and mouth appear in approximately the same location. The faces are then passed on to the pre-trained neural network to generate 128 descriptor measurements for each face. To create an RDM, we used the program’s calculated distances between the measurements for each pair of faces images. A value of zero indicates that two images are identical, and values between 0 and 1 suggest that two different images likely show the same person’s face. Values higher than 1 indicate that the two images show the faces of two different people. We found that OpenFace performed well at grouping different images of the same person’s face compared to images of different people’s faces in our image set — Supplementary Information 1 includes full 72×72 matrices showing distances between all images). To obtain a 12×12 RDM for the 12 identities, which would be comparable to the brain RDMs, we computed the mean of all cells that showed tokens of the same identity pair (Figure 2C).

#### Gabor-Jet Model

The Gabor-Jet model RDM was computed from visual descriptors of face images obtained using the Gabor-Jet model (Biederman & Kalocsai, 1997; Margalit et al., 2016; Yue et al., 2012). This model was designed to simulate response properties of cells in area V1, and has been found to correlate with psychophysical measures of facial similarity (Yue et al., 2012). In addition, Carlin and Kriegeskorte (2017) showed that the dissimilarity of response patterns to different faces in the FFA was predicted by image properties based on Gabor filters. First, we used OpenFace 2.0 (Baltrusaitis et al., 2018) to automatically detect the faces in each image, and the pictures were greyscaled. The Matlab script provided in www.geon.usc.edu/GWTgrid_simple.m was then used to create a 100 × 40 Gabor descriptor for each face. After transforming these matrices into vectors, we computed the Euclidean distance between the vectors from each pair of faces (Supplementary Information 1), and then averaged the distances across all pairs of stimuli that showed the same two identities (Figure 2C).

#### GIST Model (Faces only and whole Frames)

The Gist model RDMs were computed from visual descriptors of pictures obtained using the GIST model (Oliva and Torralba, 2001). The GIST model estimates information about the spatial envelope of scenes and it is related to perceived dimensions of naturalness, openness, roughness, expansion, and ruggedness. Weibert et al. (2018) showed that the similarity between the representations of different faces in the FFA, OFA, and posterior STS was predicted by the similarity of the different pictures computed using the GIST descriptor model. We extracted GIST descriptors both from the full picture (whole Frames) and just from the face (Faces only - we used the same stimuli as in the Gabor-Jet model). We then used the Matlab script provided in http://people.csail.mit.edu/torralba/code/spatialenvelope to compute GIST descriptors for each picture, and computed Euclidean distances between each pair of pictures (Supplementary Information 1). We finally averaged the distances across all pairs of stimuli that showed the same two identities (Figure 2C).

#### Pixel Model (Faces only and whole Frames)

Finally, we computed model RDMs based on pixel dissimilarity between each pair of pictures. Like for the GIST model, we computed this model both for the full picture (whole Frames) and just for the face (Faces only). We extracted pixel greyscale values for each image, computed Pearson correlations between the vectors of each pair of images, and used correlation distance as the output measure (1-*r*) (Supplementary Information 1). We finally averaged the distances across all pairs of stimuli that showed the same two identities (Figure 2C).

### RDMs based on Perceived properties

#### Social Trait Models: Trustworthiness, Dominance, Attractiveness, Valence, Social Traits (All)

RDMs for ratings of the 12 face identities on trustworthiness, dominance, attractiveness, and positive-negative valence were computed using Euclidean distances. For each participant and each social trait, the Euclidean distance between the ratings of each pair of identities was calculated (ratings were averaged across the two trials in which the same identity was presented), resulting in a 12×12 RDM per trait. We then averaged the matrices for the same trait across participants (Figure 2C).

We also created ‘Social Traits (All)’ RDMs combining all four social traits, by calculating the Euclidean distance between all trait ratings for each pair of identities, resulting in a 12×12 trait RDM per participant. We then computed the mean matrix for all social traits across participants (Figure 2C).

#### Perceived Similarity Model

The judgements in the Pairwise Visual Similarity Task indicated the degree of visual similarity between all possible pairs of identities. These ratings were averaged across the two trials in which each identity-pair was presented, and were reverse-coded to match the LDC and Euclidean distance measures, where a higher value indicates higher dissimilarity. The resulting values were arranged into a 12×12 face RDM for each participant and were then averaged across participants (Figure 2C).

#### Gender Model

Finally, a 12×12 RDM for gender was constructed by assigning a value of 0 to same gender identity pairs, and a value of 1 to different-gender identity pairs (Figure 2C).

Correlations between all 13 models are presented in Figure 2D.

## Statistical analysis

### Individual model analysis: RSA comparing brain RDMs to candidate model RDMs using correlation

For each individual participant and each ROI, we compared the brain RDM for faces with each of the candidate model RDMs defined above using Pearson correlation (Figure 3A). We then tested whether the correlations across participants for each ROI were significantly higher than zero, using two-sided one-sample Wilcoxon signed-rank tests (Nili et al., 2014). P-values were corrected for multiple comparisons using FDR correction (*q*=.05) across all 13 comparisons for each ROI. We also compared the correlations across all pairs of models within each ROI, in order to test which model was the best predictor of the variance in brain RDMs in each ROI. For these pairwise comparisons, we used two-sided Wilcoxon signed-rank tests and only significant FDR corrected values (for 78 comparisons) are reported.

An estimate of the noise ceiling was calculated for each ROI, in order to estimate the maximum correlation that any model could have with the brain RDMs in each ROI given the existing noise in the data. We estimated the noise ceiling using the procedures described by Nili et al. (2014). The lower bound of the noise ceiling was estimated by calculating the Pearson correlation of the brain RDM for each participant with the average brain RDM across all other participants (after z-scoring the brain RDM for each participant). The upper bound of the noise ceiling was estimated by computing the Pearson correlation of the brain RDM for each participant with the average brain RDM across all participants (after z-scoring the brain RDM for each participant). Upper noise ceilings are only shown in Supplementary Information 2.

### Weighted model-combination analysis: Weighted representational modelling

We also used weighted representational modelling (Khaligh-Razavi & Kriegeskorte, 2014; Jozwik et al., 2016; 2017) to combine individual models via reweighting and thus investigate if combinations of different model RDMs could explain more variance in representational geometries than any single model. For each combined model, we used linear non-negative least squares regression (lsqnonneg algorithm in Matlab) to estimate a weight for each component of the combined model. We fitted the weights and tested the performance of the reweighted (combined) model on non-overlapping groups of both participants and stimulus conditions within a cross-validation procedure, and used bootstrapping to estimate the distribution of the combined model’s performance (Storrs et al., 2020).

We used six different combinations of component models: *Image-computable* properties (OpenFace, GIST, GaborJet, and Pixel), *Social Traits* (comprising a weighted combination of the Trustworthiness, Dominance, Attractiveness, and Valence properties), *Perceived* properties (Trustworthiness, Dominance, Attractiveness, Valence, Perceived Similarity, and Gender), *Low-Level* properties (GIST, GaborJet, and Pixel), *High-Level* properties (Trustworthiness, Dominance, Attractiveness, Valence, Perceived Similarity, Gender, and OpenFace), and *All properties*.

Within each crossvalidation fold, data from eight participants for four stimulus identity conditions was assigned to serve as test data, and the remainder was used to fit the weights for each component of each of the six combined models. Because the crossvalidation was performed within a participant-resampling bootstrap procedure, the number of participant data RDMs present in each crossvalidation fold was sometimes smaller than eight (when a participant was not present in the bootstrap) or larger than eight (when a participant was sampled multiple times in the bootstrap). All data from the same participant was always assigned *only* to either the training or test split. A reweighting target RDM was constructed by averaging the training-split participants’ RDMs for training-split stimulus conditions, and weights were fitted to the components of each combined model to best predict this target RDM. The six resulting combined models, as well as the 13 individual models, were then correlated separately with each of the brain RDMs from test participants for test conditions, using Pearson correlation. The noise ceiling was also computed within every cross-validation fold using the same procedure as for the main analysis. In other words, we correlated (Pearson correlation) each test participant’s RDM with the average of all other test RDMs excluding their own (for the lower bound of the noise ceiling) and with the average of all test participants’ RDMs including their own (for the upper bound of the noise ceiling; upper bounds shown in Supplementary Information 6). This procedure was repeated for 30 participant crossvalidation folds within 30 stimulus-condition crossvalidation folds to provide a stabilised estimate of the noise ceiling and the performance of each model (Storrs, et al., 2020).

The cross-validation procedure was repeated for 1,000 bootstrap resamplings of participants for each face-selective ROI. From the resulting bootstrap distribution, we computed the mean estimate of the lower bound of the noise ceiling, as well as the mean of each model’s correlation with human data for both individual models and combined models (Figure 3B). Correlations between model and brain RDMs were considered significantly higher than zero if the 95% confidence interval of the bootstrap distribution did not include zero. Bonferroni correction was applied to correct for multiple comparisons. Finally, we compared each pair of models by testing whether the distributions of the differences between each pair of models contained zero. We only report pairwise differences that were significant after Bonferroni correction. Code for this analysis was adapted from here: https://github.com/tinyrobots/reweighted_model_comparison.

## Acknowledgements

This work was supported by a research grant by the Leverhulme Trust (RPG-2014-392) to LG, NK, and CM. We thank Tiana Rakotonombana, Roxanne Zamyadi, Rasanat Nawaz, and Natasha Baxter for help with stimuli preparation and with testing.

