## Supplementary Information for "FFA and OFA encode distinct types of face identity information"

#### Supplementary Information 1

##### — Matrices of image-computable models per image (i.e. before averaging across identity)

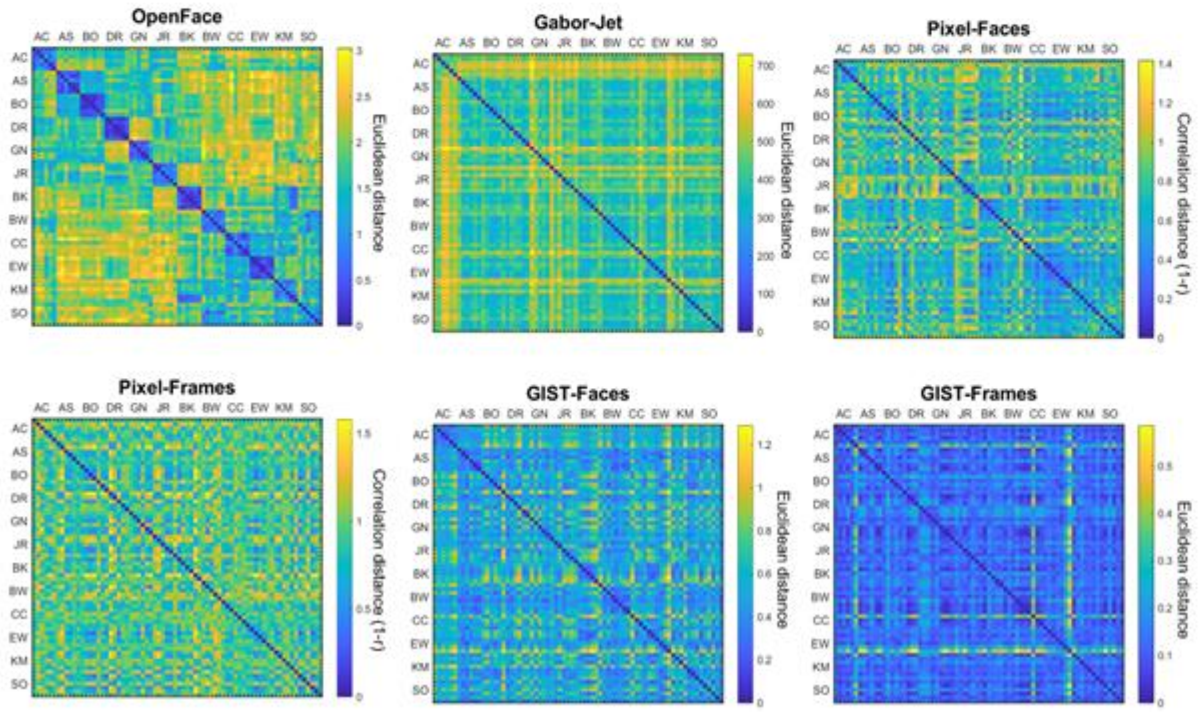


**Figure S1-1. Image-computable model representational dissimilarity matrices (RDMs) per image.** Model RDMs computed from dissimilarities between images for OpenFace, Gabor-Jet, Pixel-Faces, Pixel-Frames, GIST-Faces, and GIST-Frames. Each row/column represents a single image, and images are clustered by identity (6 images for each of the 12 identities). Each cell shows the dissimilarity between the two images in the corresponding rows and columns, with a value of zero indicating that images are identical. Matrices are symmetric around a diagonal of zeros. From these models, only the OpenFace model grouped different images of the same identity as more similar compared to images from different identities.

#### Supplementary Information 2

##### — Full results for individual model analysis

**Table S2-1**: Results of individual model analysis. For each ROI, we show the mean correlations between brain RDMs with each model, standard error (SE), Z statistics from two-sided one-sample Wilcoxon signed-rank tests, and whether correlations were significantly higher than zero. We also show the estimated lower and upper bounds of the noise ceiling for each ROI. Models are ordered by effect size.

| | Pearson correlation between RDMs | Noise ceiling |
|---|---|---|

| | | Mean *r* | *SE* | *Z* | *p* < .05 (FDR corrected) | [Lower bound Upper bound] |
|---|---|---|---|---|---|---|
| rFFA | | | | | | [0.135 0.262] |
| | Perceived Similarity | 0.109 | 0.023 | 3.689 | yes | |
| | Social Traits (All) | 0.104 | 0.031 | 2.710 | yes | |
| | Open Face | 0.101 | 0.023 | 3.461 | yes | |
| | Attractiveness | 0.090 | 0.033 | 2.687 | yes | |
| | Gender | 0.086 | 0.021 | 3.302 | yes | |
| | Valence | 0.060 | 0.023 | 2.391 | yes | |
| | Dominance | 0.058 | 0.030 | 1.640 | no | |
| | Gabor-Jet | 0.052 | 0.049 | 0.956 | no | |
| | Trustworthiness | 0.040 | 0.029 | 1.594 | no | |
| | Pixel-Faces | 0.035 | 0.044 | 0.865 | no | |
| | Pixel-Frames | 0.005 | 0.027 | 0.159 | no | |
| | GIST-Faces | -0.006 | 0.040 | 0.114 | no | |
| | Pixel-Frames | -0.018 | 0.041 | -0.478 | no | |
| rOFA | | | | | | [0.337 0.408] |
| | Pixel-Faces | 0.221 | 0.031 | 4.357 | yes | |
| | Gabor-Jet | 0.204 | 0.037 | 3.968 | yes | |
| | Pixel-Frames | 0.107 | 0.031 | 3.016 | yes | |
| | GIST-Faces | 0.104 | 0.043 | 2.216 | yes | |
| | Attractiveness | 0.092 | 0.029 | 2.843 | yes | |
| | Social Traits (All) | 0.083 | 0.031 | 1.979 | no | |
| | Gender | 0.074 | 0.021 | 2.757 | yes | |
| | OpenFace | 0.067 | 0.020 | 2.952 | yes | |
| | Dominance | 0.055 | 0.031 | 1.546 | no | |
| | Perceived Similarity | 0.039 | 0.026 | 1.416 | no | |
| | GIST-Frames | 0.025 | 0.034 | 0.746 | no | |
| | Trustworthiness | 0.011 | 0.025 | 0.400 | no | |
| | Valence | -0.016 | 0.031 | -0.573 | no | |
| rpSTS | | | | | | [0.126 0.252] |

| | | | | |
|---|---|---|---|---|
| GIST-Frames | 0.075 | 0.047 | 1.800 | no |
| Dominance | 0.052 | 0.027 | 1.800 | no |
| OpenFace | 0.040 | 0.020 | 2.129 | no |
| Social Traits (All) | 0.032 | 0.026 | 1.018 | no |
| Pixel-Frames | 0.022 | 0.030 | 0.956 | no |
| Gender | 0.020 | 0.017 | 0.956 | no |
| Trustworthiness | 0.017 | 0.032 | 0.524 | no |
| Attractiveness | 0.005 | 0.024 | 0.134 | no |
| Valence | 0.002 | 0.031 | 0.051 | no |
| Pixel-Faces | -0.003 | 0.035 | -0.113 | no |
| Perceived Similarity | -0.008 | 0.026 | -0.072 | no |
| Gabor-Jet | -0.045 | 0.040 | -1.100 | no |
| GIST-Faces | -0.048 | 0.036 | -1.368 | no |

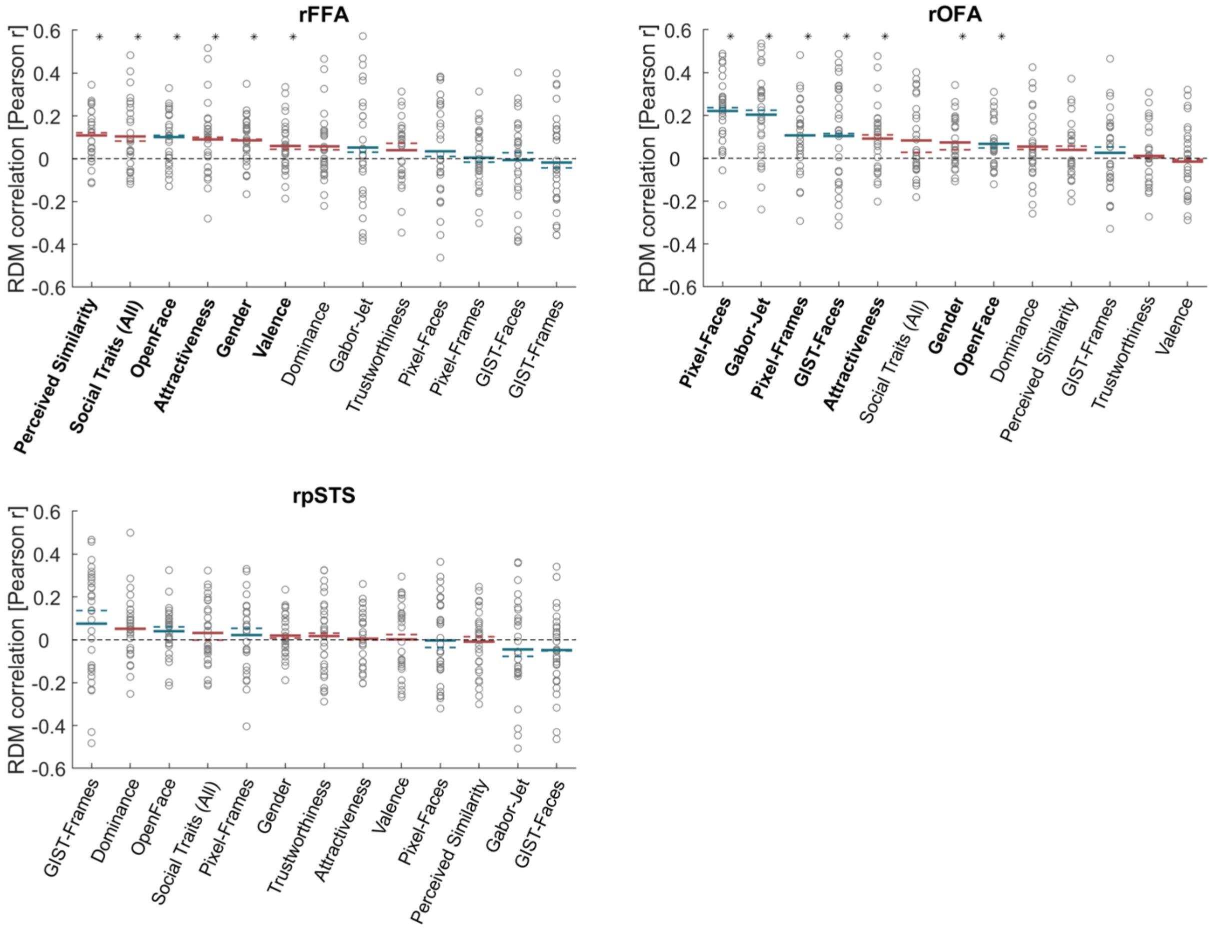

**Figure S2-1. Similarity between brain RDMs (in FFA, OFA, and pSTS) and each of the candidate models**. Circles show correlations for individual participants. Coloured lines show mean (full lines) and median (dotted lines) correlations across participants. Correlations with models based on perceived-property models are in pink, and correlations with image-computable models are in blue. Horizontal black dotted lines mark the zero correlation point. An asterisk above a bar and the name of the model in bold indicate correlations that were significantly higher than zero. Correlations with individual models are sorted from highest to lowest based on the mean correlation across participants to match the format of Figure 3 in the main manuscript.

### Supplementary Information 3

#### — Correlations between brain RDMs across different ROIs

To further test the hypothesis that the FFA and OFA encode different information about face identities, we computed correlations between each participant's brain RDM for each ROI, and the mean brain RDM for all other participants in each of the other ROIs. We predicted that if specific computations are performed in one ROI, and those computations are generalisable across participants, we should observe higher correlations between individual participants' RDMs and the average RDMs (of all other participants) for that same ROI compared to the average RDMs for other ROIs.

Figure S3-1 shows the average Pearson correlations between each participant's RDM for a specific ROI (rows) and the average of all other participants RDMs for each of the ROIs (columns). S3-1A shows correlations when using RDMs built from brain responses to faces for face-selective ROIs, and S3-1B shows correlations when using RDMs built from brain responses to voices (as control stimulus) in the same ROIs. For example, the correlation between each participant's face RDM for FFA and: (1) the average of all other participants' FFA face RDMs is 0.135, (2) the average of all other participants' OFA face RDMs is 0.114, and (3) the average of all other participants' pSTS face RDMs is 0.086. Values in the diagonal correspond to the estimates of the lower bound of noise ceiling that we presented in the main analysis. Asterisks indicate that correlations across participants were significantly higher than zero using two-sided one-sample Wilcoxon signed-rank tests.

The average correlations in each row in Figure S3-1A show that brain RDMs in each face-selective ROI are consistently more similar to other participants' RDMs for the same ROI compared to RDMs from different ROIs. These effects are specific for using RDMs based on face identities, as we do not observe these effects for FFA and OFA when using voice RDMs. We note, however, that these correlations are affected by ROI size and signal-to-noise ratio in each ROI, and that correlation values in each row are not independent from each other. Therefore, we did not carry out inferential statistics to compare correlations across ROIs.

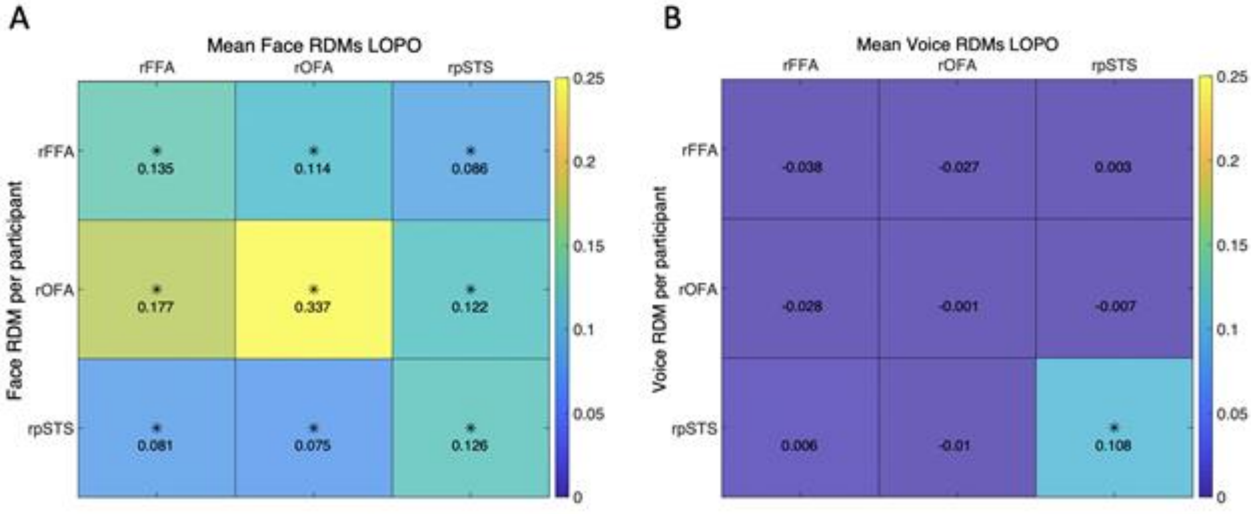


**Figure S3-1 Average (Pearson) correlations between each participant's RDM for a specific ROI (rows) and the average of all other participants' RDMs for all ROIs (columns).** LOPO: leave-one-participant-out. **A** shows correlations when using RDMs built from brain responses to faces for face-selective ROIs, and **B** shows correlations when using RDMs built from brain responses to voices in the same ROIs. Asterisks indicate that correlations across participants were significantly higher than zero.

### Supplementary Information 4

#### — Comparison of correlation measures

To investigate whether the choice of similarity measure affected the results, we repeated the main analysis, but this time using Spearman correlation and Kendall *tau-a* instead of Pearson correlations. These analyses were identical to the analysis using Pearson correlations (see Methods section), with the exception that that noise ceiling was computed after rank-transforming the RDMs (Nili et al., 2014). The pattern of results was similar across all three correlation measures (Figures S4-1 and Figure 3A). Results for correlations that were significantly greater than zero after correcting for multiple comparisons are shown in Table S4-1.

**Table S4-1: Results for individual model analysis using Spearman correlation and Kendall tau-a in rFFA and rOFA.** Results are shown only for correlations that were significantly greater than zero (after FDR correction).

| | | Spearman | | | Kendall tau-a | | |
|---|---|---|---|---|---|---|---|
| | | Mean correlation | *Z* | *p* | Mean correlation | *Z* | *p* |
| rFFA | Perceived Similarity | .11 | 3.58 | .0004 | .08 | 3.52 | .0004 |
| | Social Traits (All) | .11 | 3.10 | .0020 | .07 | 3.18 | .0015 |
| | OpenFace | .10 | 3.48 | .0005 | .07 | 3.56 | .0004 |
| | Attractiveness | .09 | 2.50 | .0123 | .06 | 2.62 | .0088 |
| | Gender | .08 | 3.07 | .0021 | .05 | 3.07 | .0021 |
| | Valence | .08 | 2.53 | .0115 | .05 | 2.71 | .0067 |

| | | | | | | | |
|---|---|---|---|---|---|---|---|
| rOFA | Pixel-Faces | .22 | 4.40 | 0.0001 | .14 | 4.30 | .0001 |
| | Gabor-Jet | .18 | 3.84 | 0.0001 | .12 | 3.88 | .0001 |
| | Pixel-Frames | .13 | 3.49 | 0.0005 | .08 | 3.38 | .0007 |
| | Gender | .07 | 2.85 | 0.0043 | .04 | 2.85 | .0043 |
| | OpenFace | .07 | 3.02 | 0.0026 | .05 | 2.89 | .0039 |
| | Attractiveness | .06 | 2.39 | 0.0169 | - | - | - |

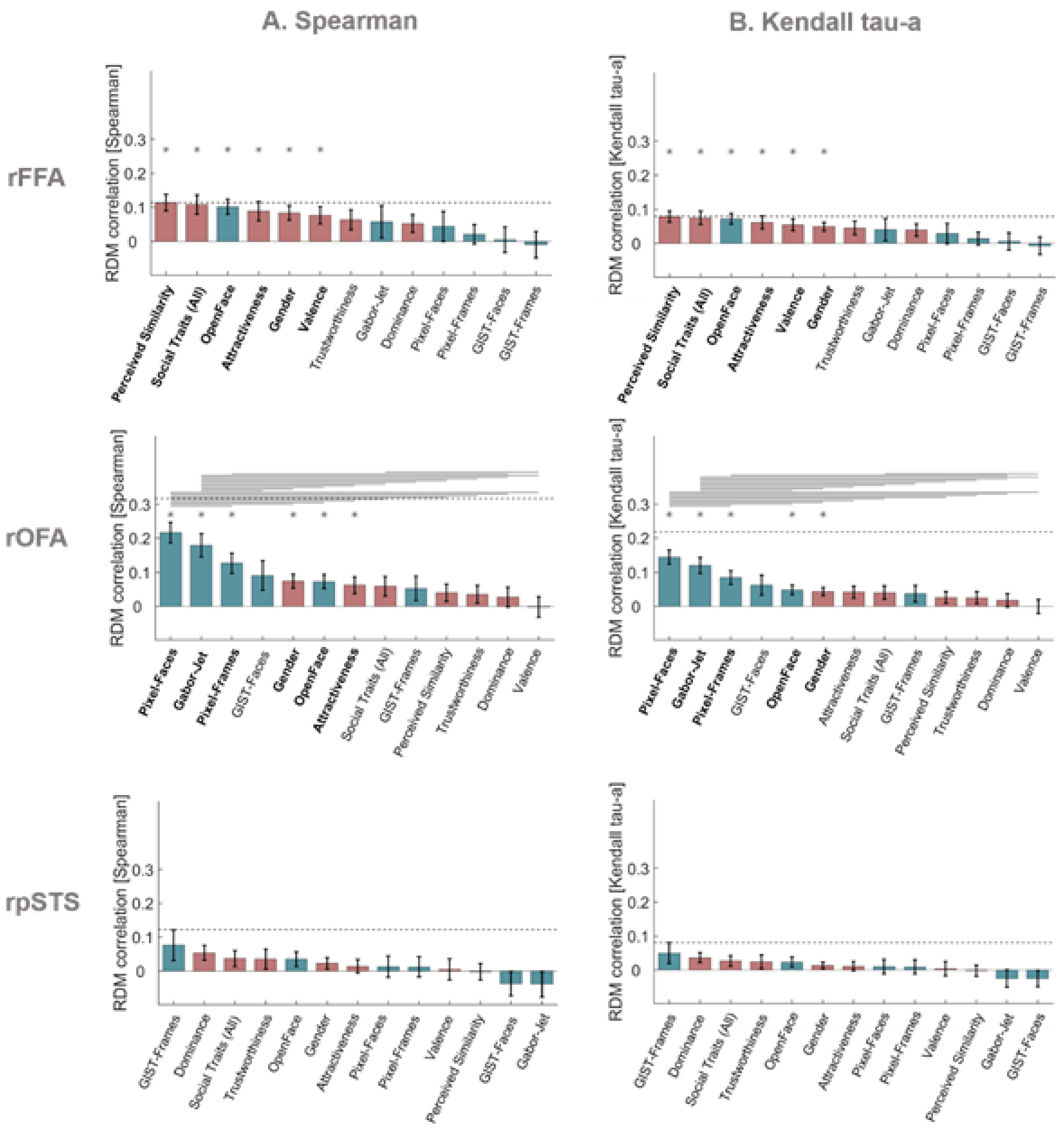


**Figure S4-1. Similarity between brain RDMs (in FFA, OFA, and pSTS) and each of the candidate models using Spearman correlation (A) and Kendall tau-a (B)**. Bars show mean correlations across participants and error bars show standard error. Correlations with models based on perceived-property models are in pink, and correlations with Image-computable models are in blue. Horizontal dashed lines show the lower bound of the noise ceiling. An asterisk above a bar and the name of the model in bold indicate correlations that were significantly higher than zero. Correlations with individual models are sorted from highest to lowest. Horizontal lines above bars show significant differences between the correlations of the two end points (FDR corrected for multiple comparisons).

### Supplementary Information 5

#### — Control analysis based on responses to voices in face-selective regions

As a control analysis, we computed representational dissimilarity matrices (RDMs) from response patterns to voices in the rFFA, rOFA, and rpSTS, and compared them with our model RDMs for faces. The voice stimuli belonged to the same 12 identities as the face stimuli and were presented interspersed among the face videos in the same runs (see Methods section). RDMs for voice identities were computed using the same procedure as for face identities (see Methods section) and were compared to model RDMs for faces using Pearson correlation. None of the correlations were significantly greater than zero after correction for multiple comparisons in the rFFA (all $p > .040$), rOFA (all $p > .103$), or rpSTS (all $p > .063$) (Figure S5-1). Pairwise comparisons showed no significant differences between the correlations of any pairs of models (all $p > .034$). The estimated lower bounds of noise ceilings were very low for rFFA ($r = -.038$) and rOFA ($r = -.001$), and higher for rpSTS ($r = .108$).

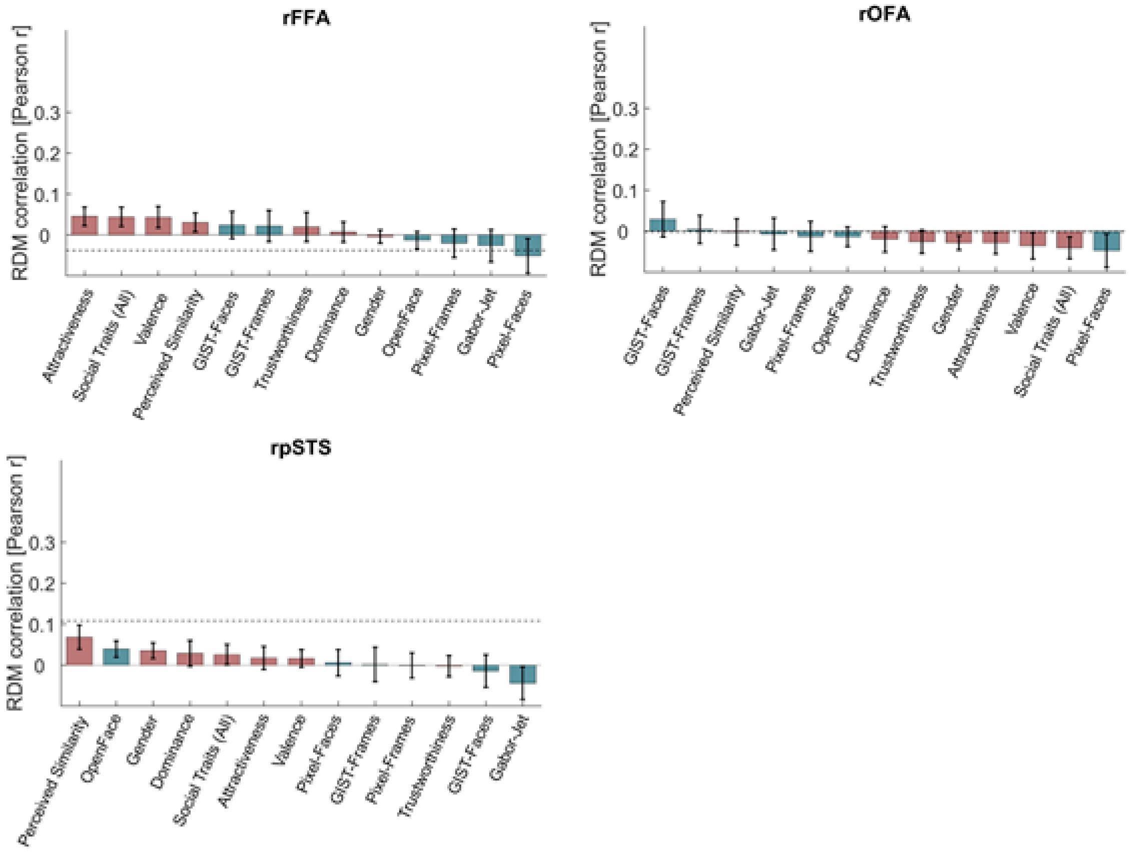


**Figure S5-1. Similarity (Pearson correlations) between brain RDMs for voices (in FFA, OFA, and pSTS) and each of the candidate models**. Bars show mean correlations across participants and error bars show standard error. Correlations with models based on perceived-property models are in pink, and correlations with image-computable models are in blue. Horizontal dashed lines show the lower bound of the noise ceiling. Correlations with individual models are sorted from highest to lowest. None of the correlations were significantly higher than zero.

### Supplementary Information 6

#### — Results of weighted representational modelling analysis

Full results of the weighted representational modelling analysis are in Table S6-1. The values in this table correspond to the results presented in Figure 3B. We also compared the correlations between all pairs of models, and the only significant difference ($p < .05$, Bonferroni corrected) was between Pixel-Faces and Valence in the OFA.

**Table S6-1: Results of weighted representational modelling analysis.** Within each ROI, we show the mean correlations between brain RDMs with each model (individual models and combined models), and whether correlations were significantly higher than zero. We also show the estimated lower and upper bounds of the noise ceiling for each ROI, and whether correlations were significantly below the noise ceiling. Models are ordered by effect size and grouped first by image-computable models, then perceived-property models, and then models that combined both types of properties. RW refers to combined and reweighted models.

| | | Pearson correlation between RDMs | | | Noise ceiling | |
|---|---|---|---|---|---|---|
| | | Mean *r* | *SE* | *p* < .05 (Bonferroni corrected) | [Lower bound Upper bound] | *p* < .05 (Bonferroni corrected) |
| rFFA | | | | | [0.089 0.286] | |
| | Open Face | 0.105 | 0.032 | yes | | no |
| | Gabor-Jet | 0.041 | 0.042 | no | | no |
| | Pixel-Faces | 0.027 | 0.040 | no | | no |
| | Pixel-Frames | 0.019 | 0.036 | no | | no |
| | GIST-Faces | 0.007 | 0.037 | no | | no |
| | GIST-Frames | -0.010 | 0.037 | no | | no |
| | RW Image-Computable | 0.063 | 0.037 | no | | no |
| | Perceived Similarity | 0.118 | 0.031 | yes | | no |
| | Social Traits (All) | 0.102 | 0.035 | yes | | no |
| | Gender | 0.094 | 0.033 | yes | | no |
| | Attractiveness | 0.091 | 0.035 | no | | no |
| | Valence | 0.059 | 0.031 | no | | no |
| | Trustworthiness | 0.049 | 0.033 | no | | no |
| | Dominance | 0.048 | 0.034 | no | | no |
| | RW Social Traits | 0.074 | 0.034 | no | | no |

| | | | | | | |
|---|---|---|---|---|---|---|
| | RW Perceived | 0.100 | 0.033 | yes | | no |
| | RW Low-Level | -0.006 | 0.035 | no | | no |
| | RW High-Level | 0.096 | 0.033 | yes | | no |
| | RW ALL | 0.086 | 0.035 | no | | no |
| rOFA | | | | | [0.237 0.372] | |
| | Pixel-Faces | 0.158 | 0.041 | yes | | no |
| | Gabor-Jet | 0.138 | 0.047 | yes | | no |
| | Pixel-Frames | 0.108 | 0.039 | no | | yes |
| | GIST-Faces | 0.087 | 0.047 | no | | no |
| | OpenFace | 0.066 | 0.041 | no | | yes |
| | GIST-Frames | 0.050 | 0.042 | no | | yes |
| | RW Image Computable | 0.089 | 0.044 | no | | no |
| | Gender | 0.082 | 0.041 | no | | no |
| | Attractiveness | 0.075 | 0.039 | no | | yes |
| | Social Traits (All) | 0.067 | 0.040 | no | | yes |
| | Perceived Similarity | 0.055 | 0.039 | no | | yes |
| | Dominance | 0.039 | 0.038 | no | | yes |
| | Trustworthiness | 0.031 | 0.040 | no | | yes |
| | Valence | -0.010 | 0.041 | no | | yes |
| | RW Social Traits | 0.037 | 0.040 | no | | yes |
| | RW Perceived | 0.033 | 0.040 | no | | yes |
| | RW Low-Level | 0.103 | 0.046 | no | | no |
| | RW High-Level | 0.019 | 0.040 | no | | yes |
| | RW ALL | 0.059 | 0.041 | no | | yes |
| rpSTS | | | | | [0.091 0.277] | |
| | GIST-Frames | 0.051 | 0.040 | no | | no |
| | OpenFace | 0.034 | 0.030 | no | | no |
| | Pixel-Faces | 0.009 | 0.034 | no | | no |
| | Pixel-Frames | 0.006 | 0.032 | no | | no |
| | GIST-Faces | -0.031 | 0.034 | no | | no |

| | | | | |
|---|---|---|---|---|
| Gabor-Jet | -0.038 | 0.037 | no | no |
| RW Image-Computable | 0.013 | 0.036 | no | no |
| Dominance | 0.054 | 0.030 | no | no |
| Social Traits (All) | 0.035 | 0.030 | no | no |
| Trustworthiness | 0.026 | 0.033 | no | no |
| Gender | 0.023 | 0.029 | no | no |
| Valence | 0.005 | 0.033 | no | no |
| Attractiveness | 0.003 | 0.029 | no | no |
| Perceived Similarity | -0.003 | 0.032 | no | no |
| RW Social Traits | 0.026 | 0.033 | no | no |
| RW Perceived | 0.031 | 0.032 | no | no |
| RW Low-Level | 0.010 | 0.038 | no | no |
| RW High-Level | 0.033 | 0.031 | no | no |
| RW ALL | 0.025 | 0.030 | no | no |

### Supplementary Information 7

#### — Individual differences and idiosyncratic representations

It could be that the lower bound of the noise ceiling and the correlations between brain and model RDMs were relatively low because of substantial individual differences in brain representations. Brain and behavioural representations of face identities could instead be idiosyncratic and thus characteristic of each individual. We considered below three ways in which we could test this hypothesis.

First, we considered the reliabilities of brain RDMs. To estimate the lower-bound of the noise ceiling, we had computed inter-subject reliabilities of brain RDMs. If, however, there are substantial individual differences in the brain RDMs, we expect that representational distances in each of the face-selective ROIs could be highly reliable within each participant but not across participants. We thus computed intra-subject reliabilities of brain RDMs by correlating the brain RDMs calculated independently from two separate testing sessions for each participant, and then averaging the correlations across participants. We note that in all other analyses in the present manuscript, the brain RDMs for each participant corresponded to the average of these two sessions. Table S7-1 shows both the inter-subject reliabilities as computed before, and the intra-subject reliabilities. For all three face-selective ROIs, we observed intra-subject reliabilities that were on average lower than the inter-subject reliabilities, suggesting that in fact, in this case, the brain RDMs are not more reliable within each individual. It is important to note, however, that there is much less data to compute intra-subject reliabilities than inter-subject reliabilities.

**Table S7-1: Inter-subject reliabilities and intra-subject reliabilities for brain RDMs.** Inter-subject reliabilities were computed as the correlation between each participant's RDM for each ROI and the mean of all other participants' RDMs for the same ROI. These correlations were then averaged across participants, and these values correspond also to our estimates of the lower bound of noise ceiling. Intra-subject reliabilities were computed as the correlation between two brain RDMs from each participant (from two separate scanning sessions, separately for each ROI). These correlations were then averaged across participants.

| | Mean inter-subject reliability | Mean intra-subject reliability |
|---|---|---|
| rFFA | 0.135 | 0.063 |
| rOFA | 0.337 | 0.079 |
| rpSTS | 0.126 | 0.094 |

Second, idiosyncratic brain representations could also result in higher correlations between each participant's brain RDM and behavioural RDMs based on their own ratings, compared to the average behavioural RDMs that we used in the main analyses. We thus repeated the main analysis using each individual's own RDMs for the rating-based perceived-property models, namely Perceived Similarity, Trustworthiness, Dominance, Attractiveness, Valence, and Social Traits (All). Figure S7-1 shows the results of this analysis. The results do not reveal higher correlations when using the individual behavioural models. In contrast, correlations with the participants' individual behavioural models are slightly lower than when using average behavioural models.

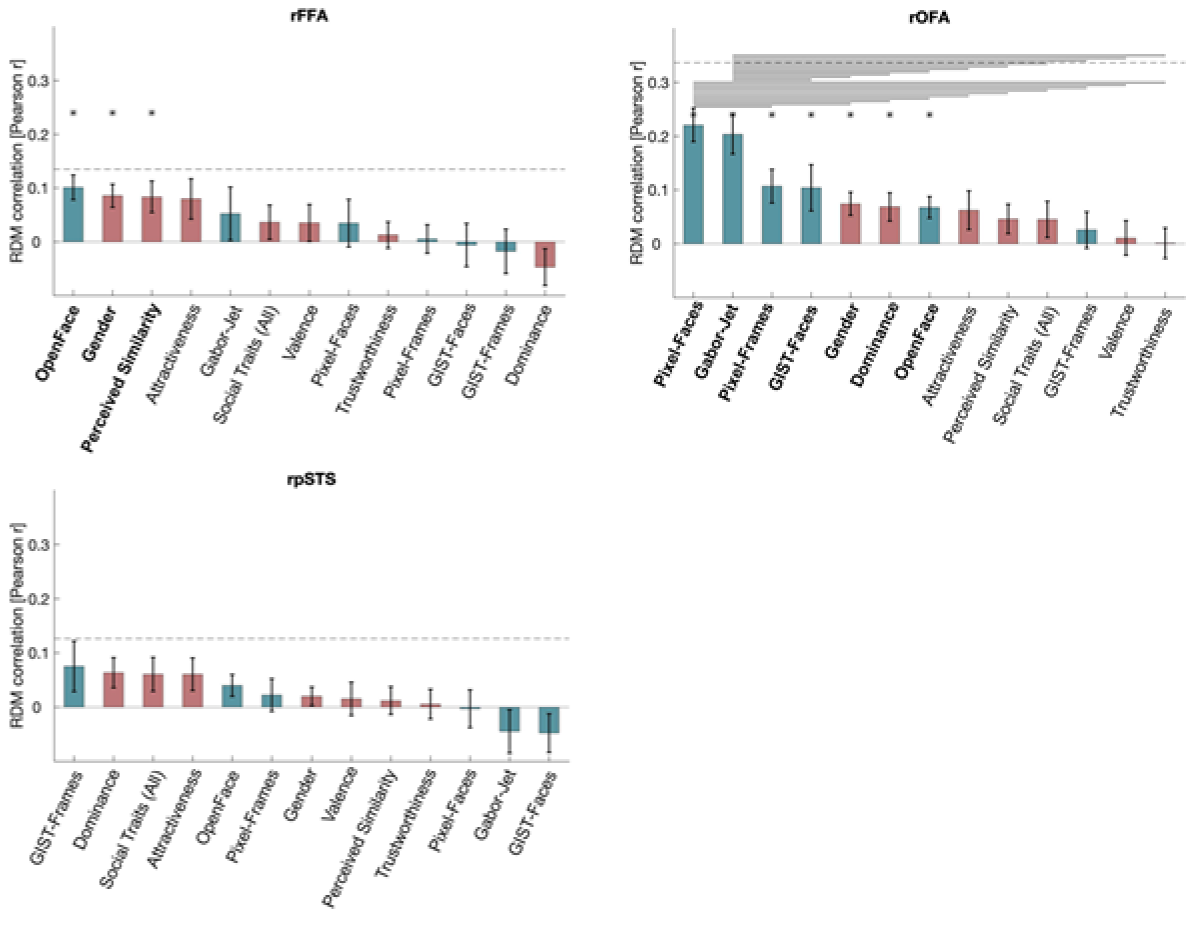

**Figure S7-1: Pearson correlations between brain RDMs (in FFA, OFA, and pSTS) and each of the individual candidate models, using behavioural models based on individual participant ratings.** Data is the same as Figure 3A, but instead of using average behavioural RDMs, each participant's brain RDM was correlated to their own behavioural RDMS for Perceived Similarity, Trustworthiness, Dominance, Attractiveness, Valence, and Social Traits (All). Bars show mean correlations across participants and error bars show standard error. Correlations with image-computable models are in blue and with perceived-property models are in pink. Horizontal dashed lines show the lower bound of the noise ceiling. An asterisk above a bar and the name of the model in bold indicate that correlations with that model were significantly higher than zero. Correlations with individual models are sorted from highest to lowest. Horizontal lines above bars show significant differences between the correlations of the two end points (FDR corrected for multiple comparisons).

A third possibility is that idiosyncratic representational geometries could result in the variance of each participant's brain RDMs being best explained by a uniquely weighted combination of candidate models (even if no set of weightings would perform well for all participants). However, we do not have sufficient data per participant to test this hypothesis here.
